# Biomarker-guided treatment strategies for ovarian cancer identified from a heterogeneous panel of patient-derived tumor xenografts

**DOI:** 10.1101/2020.01.08.898734

**Authors:** Adam C. Palmer, Deborah Plana, Hui Gao, Joshua M Korn, Guizhi Yang, John Green, Xiamei Zhang, Roberto Velazquez, Margaret E McLaughlin, David A Ruddy, Colleen Kowal, Julie Goldovitz, Caroline Bullock, Stacy Rivera, Daniel Rakiec, GiNell Elliott, Paul Fordjour, Ronald Meyer, Alice Loo, Esther Kurth, Jeffrey A Engelman, Hans Bitter, William R Sellers, Peter K Sorger, Juliet A Williams

**Author notes:** These authors contributed equally (Adam C. Palmer, Deborah Plana, Hui Gao). These authors contributed equally (Peter K Sorger, Juliet A Williams). To whom correspondence should be addressed **Corresponding authors** Juliet A Williams, PhD Novartis Institutes for Biomedical Research, 250 Massachusetts Avenue, Cambridge, MA, 02139 Peter K. Sorger, Ph.D 200 Longwood Avenue, Warren Alpert 440 Boston, MA 02115-5730, cc.

## Abstract

Advanced ovarian cancers are a leading cause of cancer-related death in women. Such cancers are currently treated with surgery and chemotherapy which is often temporarily successful but exhibits a high rate of relapse after which treatment options are few. Here we assess the responses of a panel of patient-derived ovarian cancer xenografts (PDXs) to 19 mono and combination therapies, including small molecules and antibody-drug conjugates. The PDX panel aimed to mimic the heterogeneity of disease observed in patients, and exhibited a distribution of responsiveness to standard of care chemotherapy similar to human clinical data. Three monotherapies and one drug combination were found to be active in different subsets of PDXs. By analyzing gene expression data we identified gene expression biomarkers predictive of responsiveness to each of three novel targeted therapy regimens. While no single treatment had as high a response rate as chemotherapy, nearly 90% of PDXs were eligible for and responded to at least one biomarker-guided treatment, including tumors resistant to standard chemotherapy. Biomarker frequency was similar in human patients, suggesting the possibility of a new therapeutic approach to ovarian cancer and demonstrating the potential power of PDX-based trials in broadening the reach of precision cancer medicine.

## INTRODUCTION

Epithelial ovarian cancer is among the leading causes of cancer-related death in women. The high mortality of this cancer reflects the fact that it is often at an advanced stage when diagnosed, and relatively few effective therapies are available for recurrent disease^1^. The most common treatment for advanced-stage epithelial ovarian cancer is surgical resection (if possible) followed by taxane/platinum-based chemotherapy. A high percentage of patients respond initially but the disease usually relapses and 5-year survival is around 30%^2,3^. Angiogenesis inhibitors and PARP inhibitors have improved survival^4^ though not cure rates, and overall the past decade has seen few new therapeutic strategies, emphasizing a pressing unmet medical need.

The difficulty in developing targeted therapies for epithelial ovarian cancers arises in part because the mutational spectrum of the disease does not exhibit many recurrent mutations in genes that might make attractive therapeutic targets; this is in contrast to other types of cancer for which therapies have been developed for specific, biomarker defined subtypes (e.g. lung adenocarcinoma, chronic and acute myeloid leukemia, melanoma, *HER2*^+^ breast cancer). In ovarian cancers, mutations in the P53 and homologous recombination repair pathways are common (96% and 22% of tumors respectively), but few patients have additional mutations that that can currently be used to guide therapy (reviewed in Coward et al, 2015^4^). Ovarian cancers nonetheless display substantial inter-patient heterogeneity in therapeutic response, suggesting the existence of as-yet unrecognized determinants of drug sensitivity and resistance and the potential for personalized treatment regimens. Preclinical models that reflect the heterogeneous biology of this ovarian cancer have the potential to change this situation by providing data on therapeutic vulnerabilities that can be associated with specific biomarkers.

In contrast to cell lines and conventional xenografts, patient-derived xenografts (PDXs) often preserve histopathologic and genetic features of human tumors at the time of their resection. PDX models are not an ideal mimic of human disease, in part because xenografting human tumors requires the use of mice that lack functional adaptive immune systems, but measures of drug efficacy in PDXs such as changes in tumor volume and duration of progression-free survival (PFS)) more closely resemble clinical responses than conventional xenografts or studies with cell lines. In particular, as data in this study will show, ovarian tumor PDXs recapitulate the clinically known association between BRCA loss-of-function mutations and sensitivity to the PARP inhibitor Olaparib, which has not been reproduced in panels of ovarian cancer cell lines (Supplementary Fig. S1^5–7^). PDX-based preclinical studies of patient to-patient variability in drug response have been reported for lung, breast, colon, melanoma and pancreatic cancers, including a previously described encyclopedia of >1000 PDXs^8–10^. Here we report for the first time the *in vivo* therapeutic effects of 19 different treatments, including single agents and combinations, using a panel of well-characterized ovarian PDXs from this encyclopedia. We then associated differences in response with differences in tumor gene expression, making it possible to nominate potential biomarkers for ovarian cancers. We found that a majority of ovarian PDX tumors have transcriptional or genomic biomarkers that can be used to guide the selection of treatment that elicits tumor regression as great as that associated with responses to standard of care combination chemotherapy. Whether these biomarkers will translate into humans remains unknown, but our data strongly suggest the existence of new opportunities for biomarker-guided treatment of ovarian cancer. The approach is analogous to genome-informed inhibition of oncogenes in non-small cell lung cancer^11^, but applicable to ovarian cancers and other possibly other cancers with few ‘druggable’ oncogenes by encompassing biomarkers of drug sensitivity that do not correspond to oncogenic drivers.

## METHODS

### Generation of ovarian patient-derived xenografts

Mice were maintained and handled in accordance with the Novartis Institutes for BioMedical Research (NIBR) Animal Care and Use Committee protocols and regulations. Patient tumor specimens were obtained from: the National Disease Research Interchange, Philadelphia, PA, USA; the Cooperative Human Tissue Network funded by National Cancer Institute, Rockville, MD, USA; Maine Medical Center, Portland, ME, USA. All patients were provided informed consents, samples were procured and the study was conducted under the approval of the review boards of each institution. Clinical and pathologic data were entered and maintained in appropriate databases. Generation of the PDX encyclopedia has been reported previously^9^. Briefly, tumor samples resected from ovarian cancer patients were preserved in RPMI medium and implanted in mice within 24 hours post-surgery. About 30-50mg tumor fragments were subcutaneously implanted into the flank region of athymic nude mice (Crl: Nu(NCr)-Foxn1^nu^; Charles River Laboratories). Tumor growth was monitored twice per week and successfully engrafted tumors then propagated and banked after a serial passage (p3 or p4). Flash frozen and FFPE tumor fragments were collected and used for DNA/RNA extraction and histopathological evaluation, respectively. The identity of established PDXs (defined as models propagated at least three times in mice) was confirmed by SNP48 analysis before and after PDX Clinical Trial (PCT) studies, as reported previously^9^.

### Histopathological characterization of established ovarian PDXs

Established PDXs and corresponding patient tumor samples were subjected to histopathologic examination. Freshly collected xenograft tumor samples and patient tumor samples preserved in RPMI were fixed in 10% neutral-buffered formalin for 6–24 h, processed, and paraffin embedded. FFPE sections were cut at 3.5 microns, mounted on slides, baked at 60°C for at least 30 minutes, deparaffinized and stained with hematoxylin and eosin. Images were captured using an Aperio Scanscope (Aperio Technologies).

### Genomic profiling of ovarian PDXs

RNA and DNA from ~30 mg flash frozen tumor fragments was purified using the QIAcube (Qiagen cat# 9001292) automated sample preparation platform using the Qiagen ALLPrep DNA/RNA Mini Kit (cat# 80204). The RNA concentration and integrity was evaluated with an Agilent 2100 Bioanalyzer (cat #G2940CA) utilizing the Agilent RNA 6000 Nano Kit (cat# 5067-1511) and protocol. DNA quantification was assessed with the NanoDrop 8000.

Copy number analysis was derived from profiling of total DNA on the Affymetrix genome-wide human SNP Array 6.0 chip using instrumentation and protocols from Affymetrix. The raw CEL files were returned and the QC steps were performed using Affymetrix Genotyping Console. The CEL files from chips passing QC were used for copy number analysis. Both a copy number segmentation file and a gene-level copy number value was generated using Partek Genomic Suite 6.6 genomic segmentation algorithm (Partek Inc.). CIN scores were calculated as the standard deviation of the mean copy number across chromosome arms (ArmCIN) as well as the mean of the standard deviation of copy number within each chromosome arm (FocalCIN).

Total RNA was used as input to the Illumina mRNASeq 8 Sample Prep Kit (catalog number RS-100-0801) or TruSeq RNA Sample Prep Kit v2-Sets A/B (48Rxn) (catalog number FC-122-1001 and FC-122-1002) depending on the date of the RNASeq library generation. RNA input ranged from 0.25 to 2 micrograms with RIN (RNA Integrity Number) scores from 5.1 to 10.0.

The RNASeq libraries were sequenced at a range of 75 to 100 base pair paired end reads with 7 base pair index using the standard Illumina primers. The sequence intensity files were generated on instrument using the Illumina Real Time Analysis software. The intensity files were de-multiplexed and FASTQ files created using the CASAVA software suite (version dependent upon date of analysis and current CASAVA package available – latest version used was 1.8.2).

The FASTQ files were then processed as in Korpal *et al.*^12^, modified to align simultaneously to mm10 and GRCh37 genomes and transcriptomes to allow for both human (tumor) and mouse (stroma) alignment. Mutations were called only on GRCh37. For calculating mutation rates, non-COSMIC mutations that appeared in >50 samples across a larger collection of cell line and PDX models or in dbSNP v138 were removed as likely germline variants / false positives / alignment artifacts. For TCGA, we downloaded somatic SNVs identified from whole exome sequencing and copy number alterations identified by Affymetrix SNP6.0 from the cBio portal.

### PDX Clinical Trial (PCT) and drug treatment

Treatment of established ovarian PDXs in the PCT format was described previously^9^. Briefly, ovarian PDXs between passages 4-8 were used for the PCT study, and tumors (~200 mm^3^) were enrolled on a rolling basis and treated till they reached the study end points. Maximum tolerated dose (MTD) was used for the agents that have not entered the clinics yet, while clinically relevant dose (CRD) was used for agents that are currently used or have been evaluated in the clinic. The CRD was converted from human dose by matching blood exposure **(Supplementary Table 5)**. The standard treatment schedule was 21 days. Fast growing tumors were dosed till the tumors reached ~1500 mm^3^. For slow growing tumors, treatment was maintained until tumor volume doubled at least twice. Tumor size was evaluated twice weekly by caliper measurements and the approximate volume of the mass was calculated using the formula (L’W’W) ’ (π/6), where L is the major tumor axis and W is the minor tumor axis. The duration of ‘Progression Free Survival’ (PFS) was defined as the time on therapy until tumor volume doubled. In the absence of progression or an adverse event, treatment was continued for at least 150 days. The response was determined based on the defined criteria described previously^9^: mCR, BestResponse < −95% and BestAvgResponse < −40%; mPR, BestResponse < −50% and BestAvgResponse < −20%; mSD, BestResponse < 35% and BestAvgResponse < 30%; mPD, not otherwise categorized. Mice that were sacrificed because of an adverse event before they had completed 14d on trial were removed from the data set. Objective response rate (ORR) was calculated as the percent of PDXs with partial and complete responses (mPR + mCR) to each therapy.

To correct for variability in background growth rate between PDX models, we normalized the time scale of each model to the model-specific doubling time—that is, the day at which the unabated tumor is expected to double in volume. We approximated the model doubling time as the lowest 25^th^ percentile of the doubling times among successfully engrafted tumors of that model (defined as doubling in volume and maintaining a relative tumor volume greater than –20%). We used the approximated model doubling time as a scaling factor for all measured growth curve time points within each model. Each sampled time point in the normalized growth curve for a given model was calculated as follows:

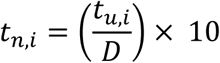

Where *t*_*n,i*_ is the *i*th time point of normalized sample *n*, *t*_*u*,*i*_ is the *i*th time point of un-normalized sample *u*, and *D* is the approximate doubling time of the model. Each normalized time point is multiplied by a factor of 10 to rescale the growth curves to approximate the original median doubling time from un-normalized growth curves. We additionally set a lower and upper limit for *D* to 5 and 20, respectively.

### Identifying drug combinations with stronger than independent action

Both PFS and tumor volume measurements were analyzed to determine if any drug combinations achieved a response better than their best constituent single-agent response, which is a standard that few combination therapies surpass *in vivo*^13^ and is evidence of additive or synergistic effect. This analysis was performed for the 9 combinations which also had response data for their constituent monotherapies. For the PFS analysis, a survival distribution was generated for the two PDX monotherapy responses that constituted a combination. To simulate the combination response expected by independent action^13^, a value was randomly sampled from each of the monotherapy response distributions, and the best of the two values was chosen. This procedure was repeated 10^6^ times to build the simulated response distribution. A number of responses equal to the number of observed combination responses was then selected at equally spaced time intervals from the full simulated response distribution. The final simulated response distribution and the observed combination response distribution were compared using a Cox proportional hazards model, with corresponding relative risk scores and p-values.

To determine statistical significance, we examined differences in responses to two PI3K inhibitors (BKM120 and CLR457) to estimate the magnitude of experimental error. A null distribution of differences in drug response was constructed by comparing tumor volume changes from BKM120 and CLR457 in 29 PDX models. For each combination therapy, volume changes expected by independent action were calculated by picking the best of the two constituent monotherapy volume changes for each PDX model. This resulting distribution of observed volume changes minus the expected independent action volume changes was compared to the null distribution (differences expected from experimental error). A Mann-Whitney test assessed whether the difference between the two volume difference distributions was significant and therefore whether the combination was significantly superior to independent action.

### Discovery of drug-response biomarkers

Identification of genes whose expression levels are predictive of drug sensitivity was performed separately for BKM120 plus binimetinib, LSU691, HKT288, and olaparib. Genes were selected that had highly variable in expression across the population of PDX tumors; specifically, those with variance in log_2_(transcript abundance in TPM) > 2. For each such high-variance gene, two metrics were calculated: (1) the significance of association (–log_10_ *P*) between transcript abundance and volume change (BestAvgResponse) by Spearman’s rank correlation, and (2) the significance (–log_10_ *P*) of the Hazard Ratio for disease progression between PDXs with less than median expression, and PDXs with greater or equal to median expression, by the Cox Proportional Hazards model. We elected not to explore different expression thresholds to avoid the risk of ‘p-hacking’. On a scatterplot of –log_10_ *P* (association with tumor volume change) versus –log_10_ *P* (association with hazard ratio for PFS), the 20 genes at the Pareto frontier were selected, that is, those most strongly associated with both volume change and PFS. For these genes, literature was searched for research articles describing the gene and the drug, or the gene and the drug’s target. Genes were only considered candidate biomarkers if literature described a mechanistically plausible interaction that might determine drug response (for example, an efflux pump with known activity against the drug). Finally, the procedure was repeated with scrambled drug response labels 100 times per drug, and the fraction of cases was counted in which any gene exhibited equal or better significance of association with tumor volume change and PFS. A gene only remained a candidate biomarker if this procedure indicated a False Discovery Rate ≤ 25% for the statistical association with drug response. Given that few candidate genes found literature support for interacting with a drug, the False Discovery Rate for both statistical association and literature support is estimated to be <5%.

## RESULTS

### Ovarian PDXs recapitulate the histopathology and genomics of human epithelial ovarian cancers

We established 49 ovarian PDXs representing a variety of histological subtypes (**Table 1**) from treatment-naïve patient tumors. H&E staining of representative samples of each subtype showed that the PDXs closely recapitulated the histopathologic characteristics of the original patient tumors (Fig. 1a) except that human stroma were replaced by mouse stroma after two passages in mice, as previously reported for other ovarian PDX studies^14–16^.

**Figure 1.**
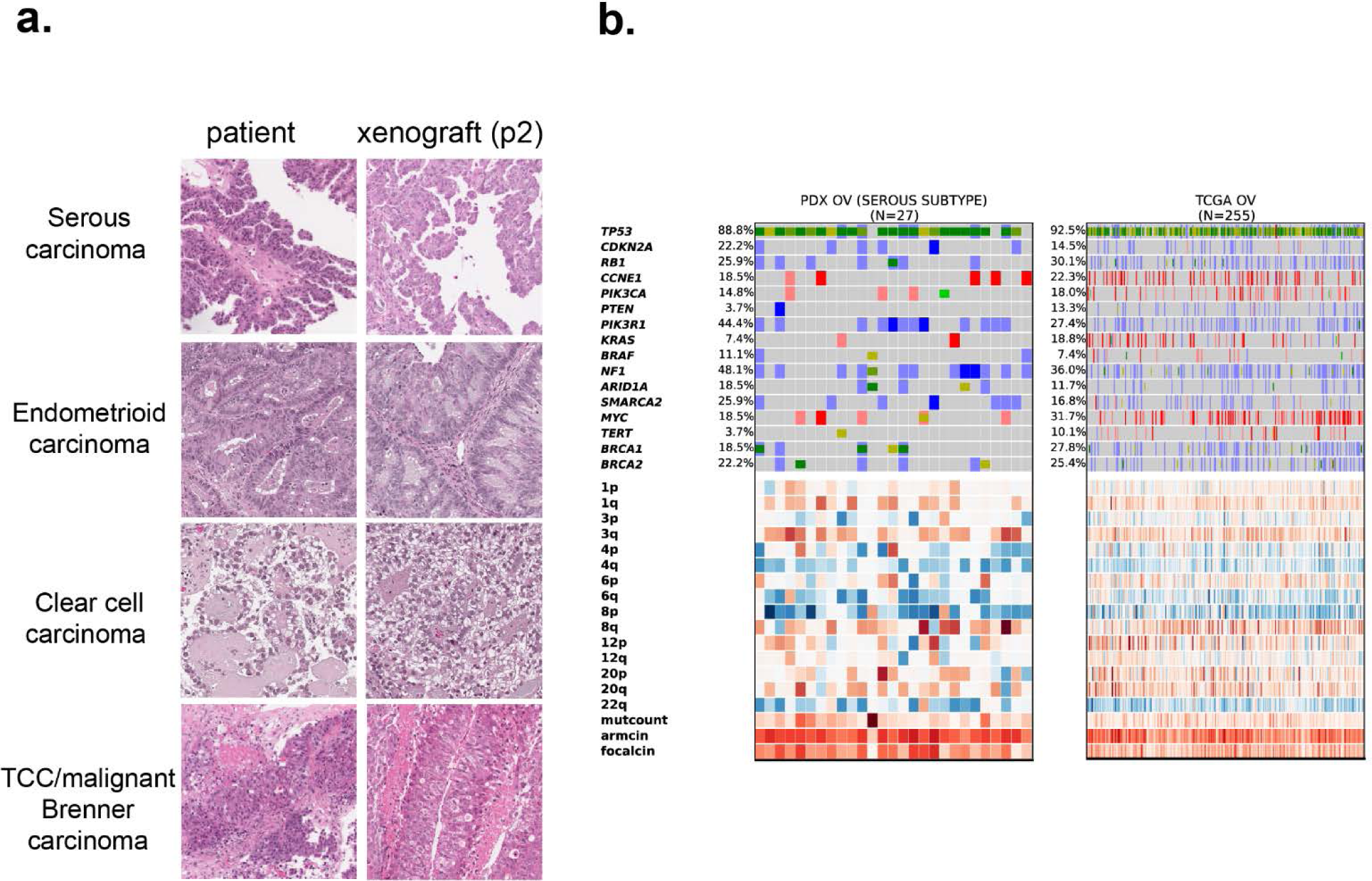
Histopathologic and genomic characterization of ovarian PDXs. **A**, Representative histologic characteristics of the original patient tumors and corresponding xenografts (passage 2) by hematoxylin and eosin staining. **B**, Genomic landscape analysis of serous ovarian carcinoma of PDXs and patient tumors (TCGA). Parenthesis, number of models per indication; blue, homozygous deletions; light blue, heterozygous deletion; salmon, amplification > 5 copies; red, amplification > 8 copies; bright green, known COSMIC gain-of-function mutations; dark green, truncating mutations / frameshift or known COSMIC loss-of-function; mustard, novel mutation. Copy number heatmap scaled from blue (deletion) to white (average CN) to red (amplification); expression heatmap scaled from blue (3 std below mean) to red (3 std above mean).

To determine if tumors in the mouse PDX cohort were representative of the diversity of human disease, we generated transcript profiles by RNA-seq and copy number profiles for 27 of the high-grade serous PDXs and compared them to data from 255 patients with high-grade serous tumors found in The Cancer Genome Atlas (TCGA)^17^ (Fig. 1b). Transcripts were mapped simultaneously to either human (tumor) or mouse (stroma). The frequency of genetic alterations across the two datasets was similar. For instance, *TP53* was mutated and/or deleted in 88% of PDXs and 92% of patient tumors, and *Cyclin E* was amplified in 19% of PDXs and 22% of patient tumors. None of 16 oncogenes or tumor suppressors often mutated in ovarian cancers demonstrated significant enrichment or depletion in high-grade serous PDX tumors relative to TCGA data. As is typical for high-grade serous ovarian cancer (and *TP53* mutant cancers in general), most PDXs and TCGA tumors demonstrated high levels of chromosomal instability and relatively low mutation rates (Fig. 1b). Genetic comparisons across all ovarian subtypes in PDXs (*n* = 38), cell lines (CCLE), and patient tumors (TCGA) is available in Supplementary Fig. S2 and **Supplementary Table 1.** We conclude that our patient-derived tumor xenograft population is a representative sample of the genetic variability of human ovarian tumors.

### Testing new therapies for ovarian cancer in PDX libraries

To identify new therapeutic strategies for advanced-stage epithelial ovarian cancer, 21 therapeutic approaches involving 10 single agents and 11 drug combinations (**Supplementary Tables 2, 3**) were screened across 27 high-grade ovarian PDXs (18 serous, 5 mixed, 2 NS, 1 clear cell, 1 endometrioid) and two additional gynecologic PDXs with unconfirmed origin. Maximum tolerated dose was used for agents that have not entered the clinics, while clinically relevant dose was used for agents that are currently used or have been evaluated in humans (Methods, **Supplementary Table 5**). As reported previously^9^, good reproducibility is observed when the same PDX tumor is challenged with the same drug in different mice. In a data set comprising 440 examples of replicate treatment for a single type of PDX (2138 animals total) fewer than 10% of responses differed from the consensus RECIST response by more than 1 category (categories comprised: CR, complete response; PR, partial response; SD stable disease; or PD, progressive disease). Additionally, the distinction between any response (CR, PR, SD) and no response (PD) was consistent in 95% of individual animals. This finding justified the use of one animal per tumor per treatment (a 1×1×1 design) to screen multiple treatment options in a population of many tumors.

We use RECIST criteria only to summarize response rates in a manner comparable to human trials, while for all analyses we used quantitative measures: change in tumor volume, and duration of progression-free survival (PFS; defined as time until tumor volume reached 200% of baseline^6^). Treatments were maintained for up to 150 days unless animal welfare considerations required euthanasia. Combination chemotherapy (carboplatin/paclitaxel at clinically relevant doses) served as a standard-of-care comparator and elicited response rates comparable to those observed in patients with high-grade serous carcinoma (du Bois *et al*, 2003): objective response rate (ORR, which is the sum of the CR and PR rates) was 62% in PDXs vs. 67% in humans, and disease control rate (the sum of CR, PR, and SD) was 81% in PDXs vs. 90% in humans. We found that all drugs and drug combinations tested were well-tolerated, with no significant drug-induced weight loss.

The drug sensitivity landscape was heterogeneous: no single therapy had a better response rate than carboplatin/paclitaxel (62%), but 90% of tumors exhibited an objective response to at least one treatment (Fig. 2, **Supplementary Tables 1, 2, 3**). Change in tumor size was strongly correlated with duration of PFS (Spearman correlation −0.77, *P* = 10^−109^). A number of novel combination therapies showed promising activity; in nine cases it was possible to compare the activity of a combination with its constituent drugs administered individually to determine whether its activity surpassed independent action (in which bet hedging allows populations to benefit from the best of multiple monotherapy activities^13^). Activity surpassing independence is evidence of drug synergy or additivity. When each individual tumor’s response to a combination therapy was compared to the best of its responses to the single drugs in that combination, changes in tumor volume and PFS were no better than monotherapy responses for eight of nine combinations, except for a combination of the PI3K inhibitor BKM120 and MEK inhibitor binimetinib (**Supplementary Table 4**). Notably, three PI3K inhibitors (BKM120, BYL719, and CLR457) when used as monotherapies resulted in modest and short-lived responses (by tumor volume and PFS respectively), as previously observed in human ovarian cancer^18^. In contrast, BKM120 plus binimetinib (Fig. 3a) was superior to either drug alone and to a model of independent action with respect to both changes in tumor volume (Fig. 3c) and duration of progression-free survival (Fig. 3b). The BKM120 plus binimetinib combination achieved 48% ORR as compared to 21% for BKM120 and 4% for binimetinib, and a median PFS >100 days as compared to 43 or 14 days. For the combination vs. BKM120 alone, the hazard ratio (HR) was 0.24 (*P*=0.01 log-rank test) and for the combination vs. binimetinib alone, the Hazard Ratio (HR)=0.09 (*P*=10^−5^); relative to an independent action model, the combination achieved an HR=0.27 (*P*=0.01) demonstrating highly superior activity (**Supplementary Table 3**). Synergy between PI3K and MEK inhibitors in PDX tumors is consistent with extensive evidence that the MAPK and PI3K pathways interact and that MAPK activation can mediate resistance to PI3K inhibitors^18^.

**Figure 2.**
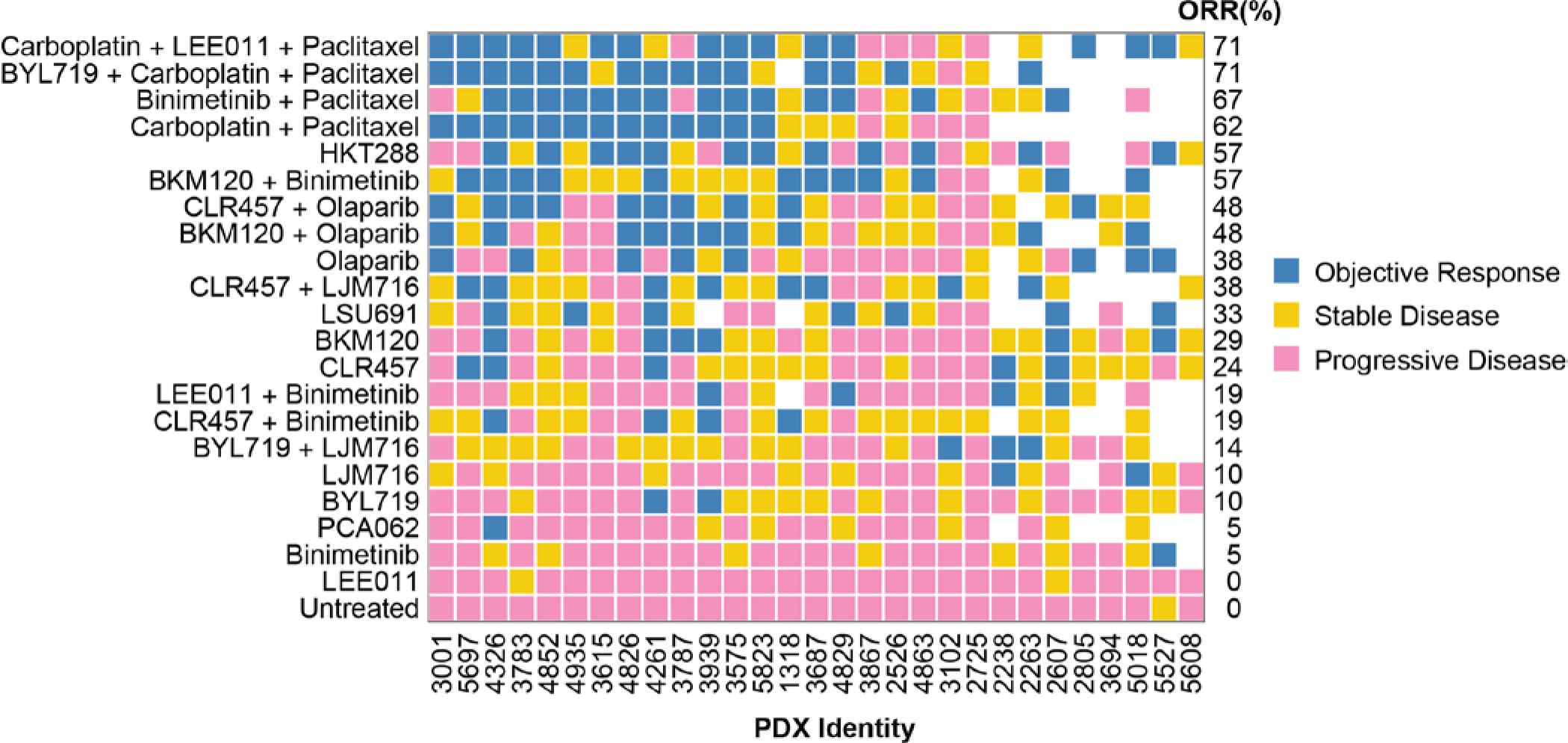
Tumor responses to novel treatments and standard chemotherapy in PDX-based clinical trials. 21 treatments were tested in 29 PDXs; each square represents a treated PDX. Objective response rate (ORR) is the percentage of PDXs with partial and complete responses to each therapy.

**Figure 3:**
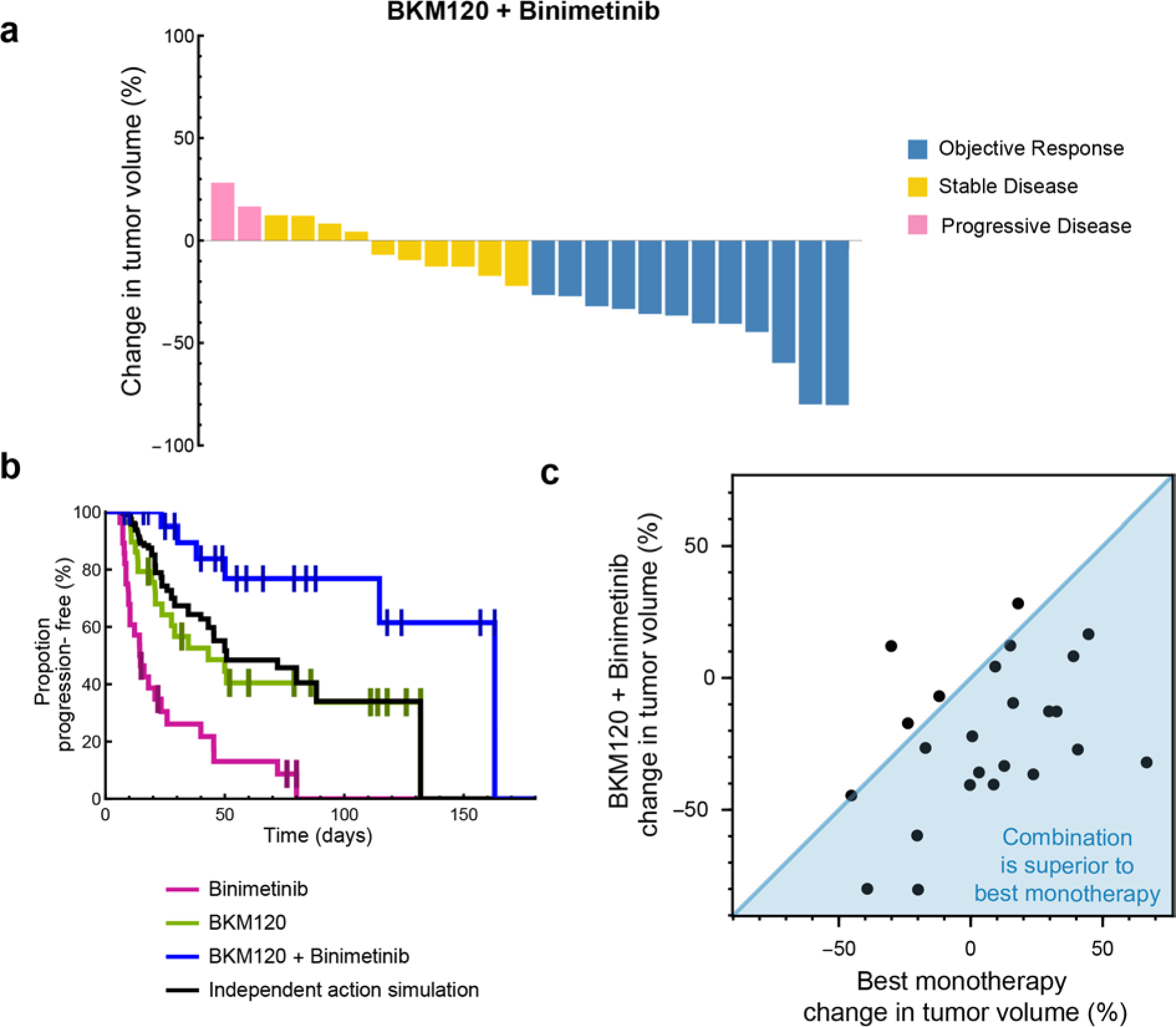
BKM 120 plus Binimetinib combination is superior to response expected by independent action. **A**, Tumor volume changes in ovarian PDXs (n=24) treated with BKM120 + Binimetinib. Color indicates RECIST response. **B**, Kaplan-Meier PFS curve of PDXs treated with: BKM120 (n=29), Binimetinib (n=28), and BKM120 + Binimetinib (n=24). Responses from each monotherapy arm were randomly sampled, and the best of the two responses was used to simulate the benefit of BKM120 + Binimetinib expected due to independent action alone. **C**, BKM120+Binimetinib tumor volume change per PDX plotted against best monotherapy volume shrinkage (BKM120 or Binimetinib response) for the same tumor (n=23).

Three drugs elicited objective responses in 25% or more of ovarian PDXs as single agents: (i) HKT288, an antibody-drug conjugate directed against Cadherin 6 (CDH6) (whose development we recently described^19^), (ii) LSU691, a small molecule inhibitor of Nicotinamide Phosphoribosyltransferase (NAMPT), and (iii) Olaparib, a small molecule inhibitor of Poly-ADP-Ribose Polymerases (PARP). In PDX trials it is possible to compare directly the response of a single tumor to different drugs and this showed that HKT288, LSU691, and BKM120 plus binimetinib induced responses in different sets of tumors, including those that were resistant to chemotherapy (Fig. 2). As a result, 88% of tumors responded to either HKT288, LSU691, olaparib, or BKM120/binimetinib in contrast to a 62% response rate for carboplatin/paclitaxel chemotherapy. While our data suggest these targeted therapies may collectively be active against a larger proportion of tumors than standard chemotherapy, for this observation to be clinically useful a means to guide drug choice is required.

### Identifying biomarkers for targeted therapies using PDX libraries

To identify predictors of drug response we compared treatment data for PDX models with RNAseq profiles. We limited the search to ~5000 protein-coding genes having a high variance in transcript abundance between PDX models. For each gene we calculated the significance of the correlation between transcript expression and treatment-induced changes in tumor volume change, and the significance of response duration (as assessed by hazard ratio) between tumors with higher or lower than median expression (Methods). We considered a gene a candidate response biomarker if it satisfied three criteria: (1) the gene was at the ‘Pareto front’ of significance - that is, among the 20 most significant in correlating with volume change *and* response duration; (2) the statistical association had a false discovery rate below 25%, based on simulations in which drug responses were scrambled among tumors; (3) a literature search revealed a mechanistic relationship between the gene product and the drug or its target (Methods).

This procedure revealed candidate biomarkers for all four targeted therapies. Resistance to BKM120 plus binimetinib was predicted by high expression of the multi-drug efflux pump *ABCG2* (Fig. 4a). Binimetinib and similar MEK inhibitors are known to be substrates of *ABCG2*^20^ and we found that the association with resistance was independently supported by binimetinib responsiveness in a panel of breast cancer PDXs^21^ (correlation of *ABCG2* expression with volume change ρ=0.33, *P*=0.046, n=37; proportional hazard of low vs high *ABCG2* groups HR=0.50, *P*=0.08) (Fig. 4b). Sensitivity to the NAMPT inhibitor LSU691 was predicted by high expression of *HCLS1*, an anti-apoptotic protein that is activated by the NAMPT/NAD^+^/SIRT1 pathway. This interaction has previously been proposed as a therapeutic target in leukemia^22^ and our data suggests that it also affects responses to NAMPT inhibition in ovarian tumors. Sensitivity to the CDH6-targeting antibody HKT288 is predicted by high expression of CDH6 itself, as was previously reported based on CDH6 immunohistochemistry^19^. Finally, the best predictor of olaparib response is loss of *BRCA1* and/or *BRCA2* function (defined here as mutation of *BRCA1* or *BRCA2*, or silencing of the *BRCA1* transcript^23^) (Fig. 4). *BRCA1/2* loss is a clinically validated biomarker of olaparib response^24^ and reproducing this association serves as a validation of our PDX-based approach. In contrast, our analysis of cancer cell line studies finds they contain no such association (Supplementary Fig. S1). Our search for predictive transcripts identified *USP51* expression as being similarly predictive as *BRCA* loss-of-function; *USP51* is recruited to double-strand breaks and regulates responses to DNA damage, including the assembly and disassembly of BRCA1 foci^25^. These data suggest that a simple test involving measuring the levels of four transcripts might be sufficient to identify tumors with a higher than average rates of response to four targeted therapies. Moreover, when PDX tumors were divided into ‘treatment-eligible’ or ‘treatment ineligible’ groups based on these biomarkers (*BRCA* loss-of-function was used for olaparib), response rates were high in eligible PDX tumors (ORR 58% to 77%) and low in ineligible tumors (ORR 0% to 11%) (Fig. 4b).

**Figure 4:**
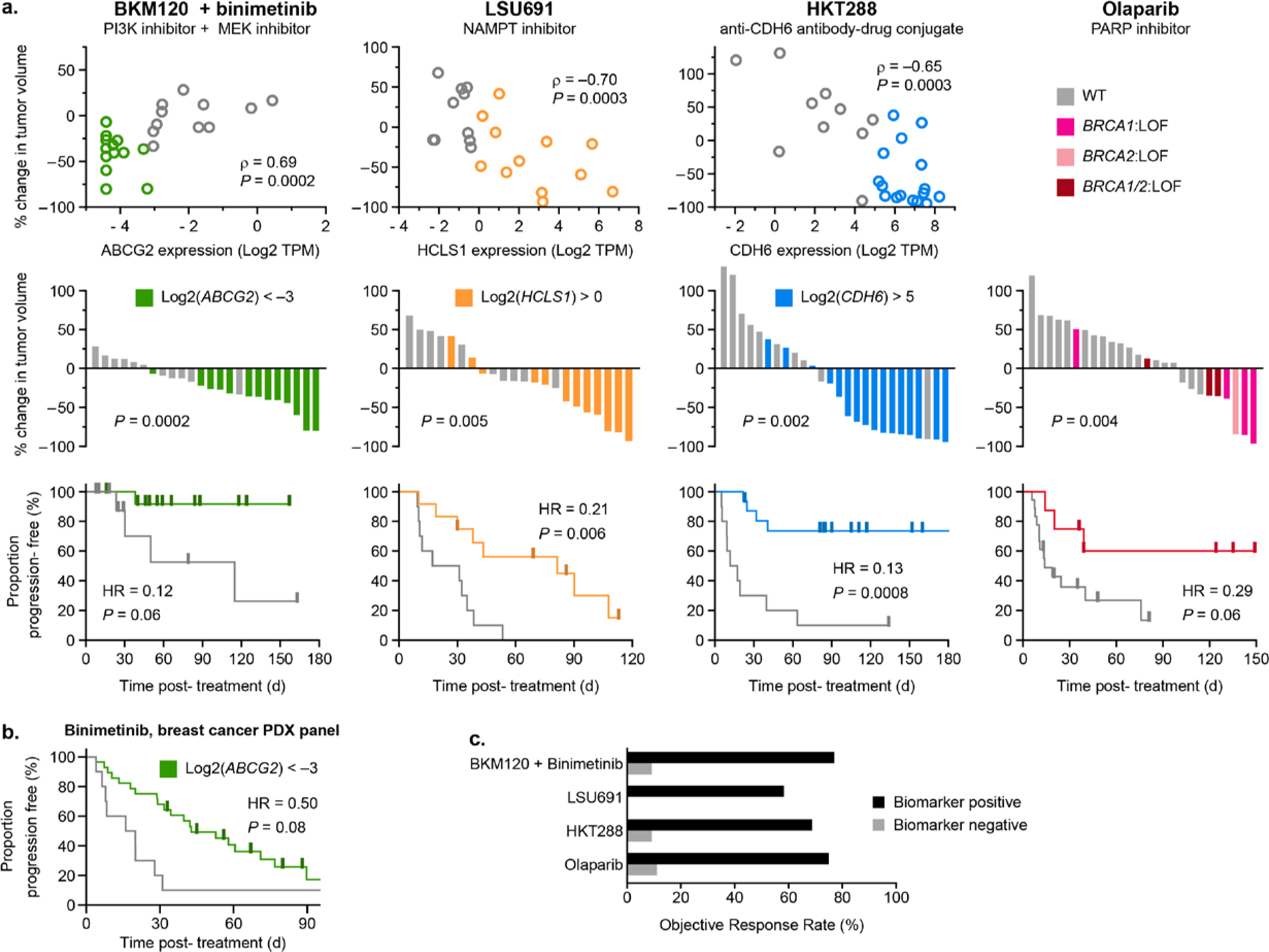
Discovery of novel biomarkers for targeted therapies in ovarian cancer. **A**, PDX treatment volume changes plotted against that model’s biomarker expression (TPM= transcripts per million) and as waterfall plots. Kaplan-Meier PFS curves are stratified into biomarker negative (gray) or positive (colored) groups (total n=24, 22, 26, and 26 respectively). **B**, Validation of *ABCG2* biomarker in a separate data set measuring duration of binimetinib response in breast cancer PDXs (n=39). **C**, Objective response rate among biomarker positive and negative indications for each therapy.

We then used TCGA primary tumor transcriptional and survival data to test whether these four biomarkers are associated with the rate of disease progression. When we compared the proportion of disease-free patients in the biomarker positive and negative groups for *ABCG2*, *HCLS1*, *CDH6*, and *BRCA1/2*, no biomarker was predictive of disease-free duration (Supplementary Fig. S3). Thus, the biomarkers appear to be predictive of drug responses, not the rapidity of disease progression.

### Most ovarian PDXs are candidates for biomarker-guided treatments

A large majority of PDX tumors (89%) were biomarker-positive candidates for one or more of four therapies: BKM120 plus binimetinib, LSU691, HKT288, or olaparib (Fig. 5a). Analysis of the same biomarkers in human primary epithelial ovarian cancers (n= 232) in The Cancer Genome Atlas revealed that a similar fraction of human tumors (93%) are positive for one or more of the same biomarkers. Furthermore, the proportion of biomarker positive tumors in TCGA did not significantly differ between rapidly progressing disease (less than 12 months disease-free) and better controlled disease (greater than 12 months disease-free), suggesting that biomarker-guided targeted therapies may be options in tumors responsive to chemotherapy as well as primary progressive disease (Fig. 5a). Among PDX tumors, 19% were ineligible or did not respond to a biomarker-indicated treatment, 33% experienced disease control from one indicated treatment, and 48% had disease control from two or more indicated treatments. When each PDX tumor was assigned its one best biomarker-indicated treatment response (by tumor volume change), the resulting set of responses was comparable to the highly effective treatment of carboplatin/paclitaxel, both by volume change (Fig. 5b. 5c; no significant difference by Mann-Whitney test, *P*=0.17) and by response duration (Fig. 5d; no significant difference in hazard ratio, *P*=0.44). Thus, a majority of ovarian PDX tumors respond strongly to a biomarker-guided treatment (ORR 88% in eligible tumors). Importantly, among the 38% of tumors that did not respond to paclitaxel/carboplatin, most (5 of 8) had a predictable objective response to one of these targeted therapies (Fig. 5e), further supporting the hypothesis that biomarker-guided therapies may be an effective option for chemotherapy-resistant ovarian cancers.

**Figure 5:**
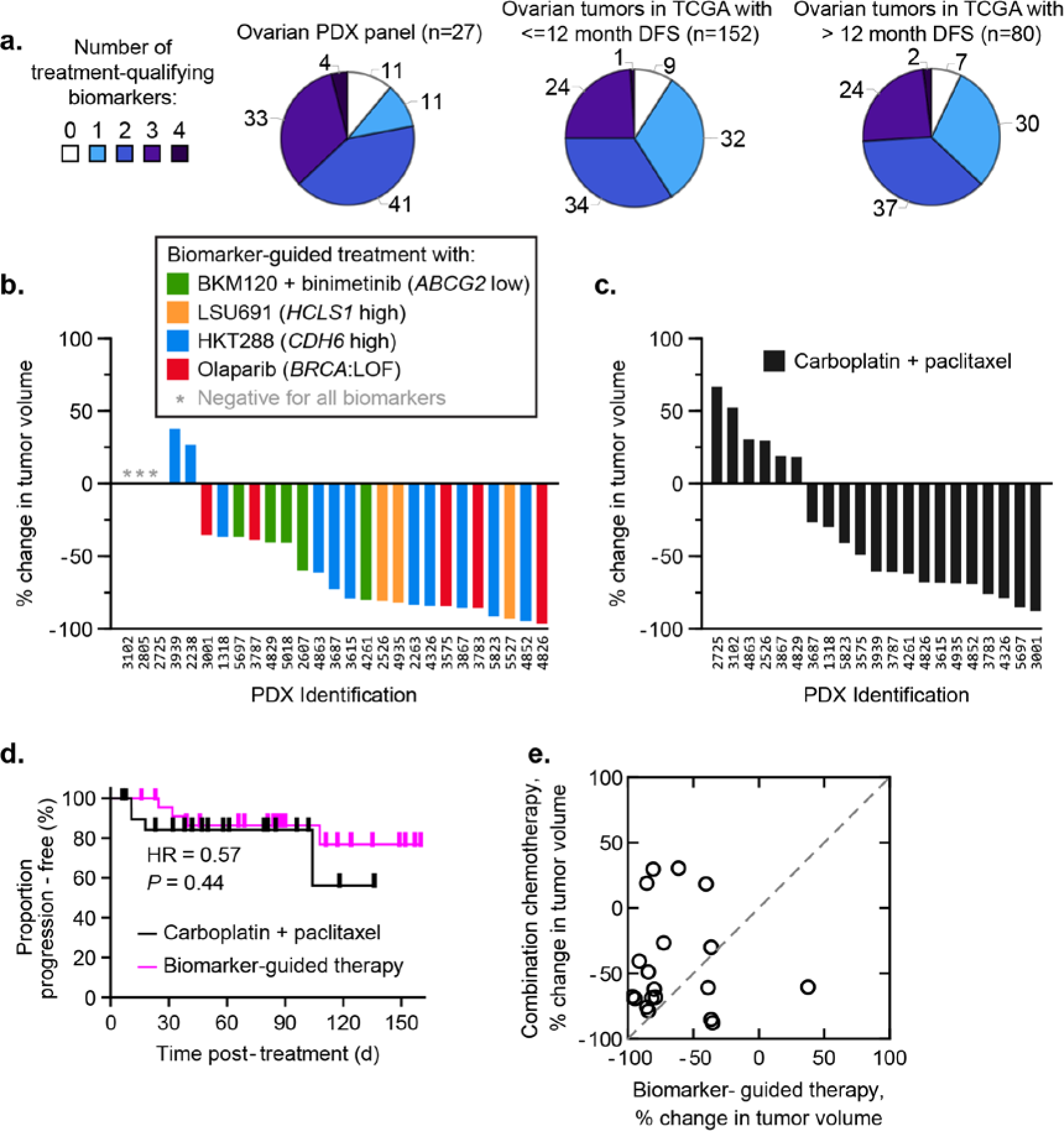
Most ovarian tumors respond to a biomarker-guided treatment. **A**, Proportion of ovarian tumors possessing one or more treatment-qualifying biomarkers (for BKM120 + binimetinib, LSU691, HKT288 or Olaparib). **B**, Tumor volume changes in ovarian PDXs each treated with their best biomarker-guided therapy. **C**, Waterfall plots of PDX responses to standard of care chemotherapy (carboplatin + paclitaxel). **D**, Kaplan-Meier PFS curves of PDXs treated with biomarker-guided therapy (n= 24) as compared to standard of care paclitaxel/carboplatin (n=21). **E**, Biomarker-guided therapy tumor volume change per PDX plotted against combination chemotherapy volume shrinkage (for the same animal; n=19).

## DISCUSSION

Approvals of the anti-angiogenesis inhibitor bevacizumab and PARP inhibitors have contributed to improved survival for ovarian cancer patients, however, the path to further progress has been unclear in the context of a difficult-to-target mutational landscape^26^. Moreover, standard combination chemotherapy is sufficiently effective to provide a high bar to a new therapy while nonetheless being ineffective for some patients and ultimately inadequate with respect to lasting control. The panel of patient-derived ovarian tumor xenografts described here was established to enable empirical comparison of multiple drug and drug combinations against chemotherapy at the level of individual human tumors. The PDX models were established with the goal of capturing the clinically observed histological heterogeneity and mutational spectrum of human ovarian cancer. At present systemic chemotherapy for epithelial ovarian cancer is consistent across multiple histological subtypes, making it appropriate to study these subtypes together^4^. These PDXs as well as their genomic sequences are freely available to the research community through the PRoXe website (www.proxe.org)^27^.

Strong anti-tumor activity (greater than 25% ORR) was observed for three single agents and one synergistic drug combination, but as is common for targeted therapies, each drug or combination was active in only a subset of tumors. We identified genes whose expression was significantly associated with drug sensitivity for each of the four therapies (for olaparib, *BRCA* status is a established biomarker^24^), making it possible to stratify tumors into likely responders and non-responders. For example, a combination of the PI3K inhibitor BKM120 plus the MEK inhibitor binimetinib was most active in PDX tumors with low expression of the *ABCG2* efflux pump, which has previously been shown to export binimetinib^20^; we confimed that *ABCG2* also predicts binimetinib response in breast cancer xenografts (Fig. 4b). The performance of this biomarker, and others described in this study, was similar to that of *BRCA* loss-of-function mutations in predicting olaparib sensitivity, which is a clinical test in breast and ovarian cancers^24,28^. In PDX models, four treatment options exhibited unstratified response rates between 31% and 48%, and choosing among these using biomarkers achieved an aggregate objective response rate of 88%. Thus nearly all PDX tumors tested had the potential to benefit from a biomarker-guided precision therapy approach.

Our data suggest that biomarkers other than oncogenic mutations or amplifications can be used to predict the effectiveness of a range of targeted therapies (e.g. antibody-drug conjugate, kinase inhibitor, inhibitor of a metabolic enzyme, etc); it is particularly helpful when these molecular features have a logical or mechanistic connection to drug response. In the case of ovarian cancer and the therapies studied here, analysis of biomarker prevalence in TCGA suggests that a precision medicine approach might be applicable to > 90% of patients. We note however, that the retrospective approach we used to identify predictive biomarkers does not involve verification in a second independent set of tumors (except in the case of *ABCG2* and binimetinib^21^). In addition, no pre-clinical study of a therapy is a guarantee of safety and efficacy in humans. Thus, the biomarkers described in this paper primarily constitute a proof of concept for precision therapy in ovarian cancer rather than an actionable set of predictions.

Possible clinical uses of biomarker-guided treatments for ovarian cancer include single-agent treatments for relapsed or chemotherapy-resistant disease, maintenance therapies following an initial course of chemotherapy (as PARP inhibitors are currently used ^29^), or in combination with chemotherapy. The present studies do not investigate responsiveness in the setting of recurrent disease, whose high incidence is a challenge in treating ovarian cancers. The likelihood of cross-resistance among targeted therapies and chemotherapy is also unclear, although at the population-level chemotherapy activity is correlated with olaparib activity, suggesting some cross-resistance, but not with other targeted therapies (HKT288, BKM120/binimetinib, LSU691), suggesting little cross-resistance in those cases. All of these questions can be subjected to pre-clinical investigation using additional PDX cohorts. Clinical testing of such a strategy, which is not limited to the therapies studied here, would most efficiently be conducted using a master protocol in which multiple biomarker-guided therapies are evaluated, as with the Lung-MAP trial^30^.

PDX drug trials have important limitations but they represent a unique setting in which to answer scientific questions that cannot be addressed in humans. Among the limitations of PDXs are potential biases introduced by low engraftment rates, by the replacement of human stroma with mouse stroma, or by smaller sizes of tumors in mice (particularly as compared to human ovarian cancers) and attendant limits on intra-tumor heterogeneity. The tumors used in this study were donated anonymously and thus, it is neither possible nor ethical to compare responses in animals to patient histories. Reassuringly, however, the population of PDX tumors exhibited the same response rate to first-line chemotherapy as do patients with ovarian cancer, and they reproduce *BRCA* loss-of-function as a predictor of olaparib response. Key advantages of PDX trials include the ability to determine how a single tumor compares to a range of therapies and to thereby gain insight into biomarkers that are predictive of drug response rather than prognostic. Ultimately, these data demonstrate the capacity of population-based PDX trials to support biomarker discovery and inform the design of future clinical trials and patient selection strategies. These capabilities are today accessible to academic researchers through consortia, notably PDXNet^31^ and the Public Repository of Xenografts^27^.

## ACKNOWLEDGEMENTS

We thank A. Färkkilä for helpful discussions and comments on this manuscript.

## SUPPLEMENTARY DATA

**Supplementary Fig. S1:**
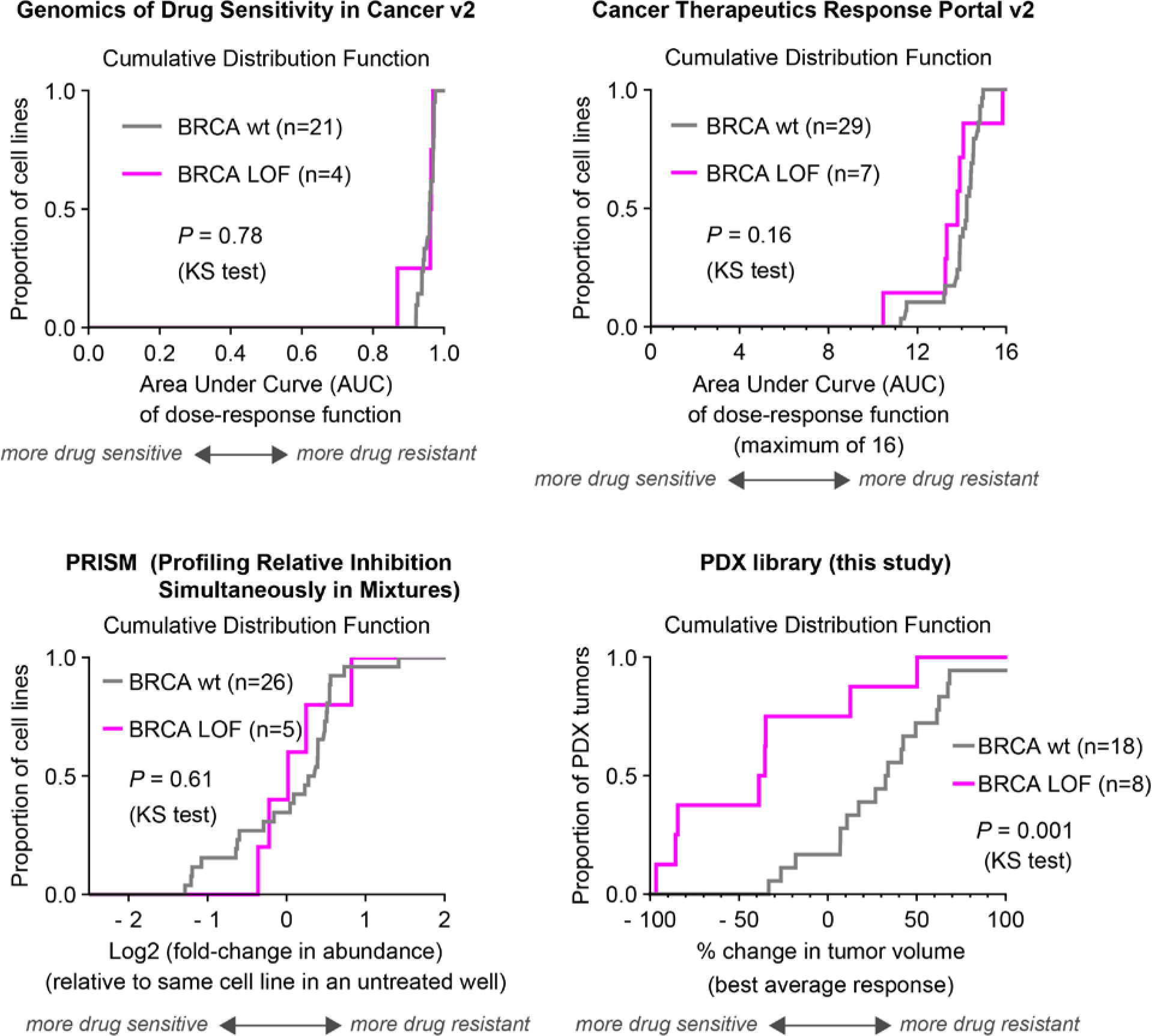
Ovarian cancer cell lines do not reproduce the clinical association between BRCA loss of function and sensitivity to olaparib, but patient-derived ovarian tumor xenografts do. Measurements of olaparib response in ovarian cancers were obtained from systematic screens of drug response in cancer cell lines (Genomics of Drug Sensitivity in Cancer v2^5^, Cancer Therapeutics Response Portal v2^6^, and PRISM^7^). Ovarian cancer cell lines were sorted into two groups: *BRCA* loss-of-function (inactivating mutation in either *BRCA1* or *BRCA2*) or *BRCA* wildtype. Differences in olaparib sensitivity between groups were illustrated by plotting cumulative distribution functions of each group’s response metric, and tested for significance with the Kolmogorov-Smirnov test. The same analysis was also applied to olaparib response in patient-derived tumor xenografts, as reported in this study. Sample sizes were comparable between studies, but only patient-derived ovarian tumor xenografts demonstrated an association between *BRCA* loss-of-function and sensitivity to olaparib (*P* = 0.001).

**Supplementary Fig. S2.**
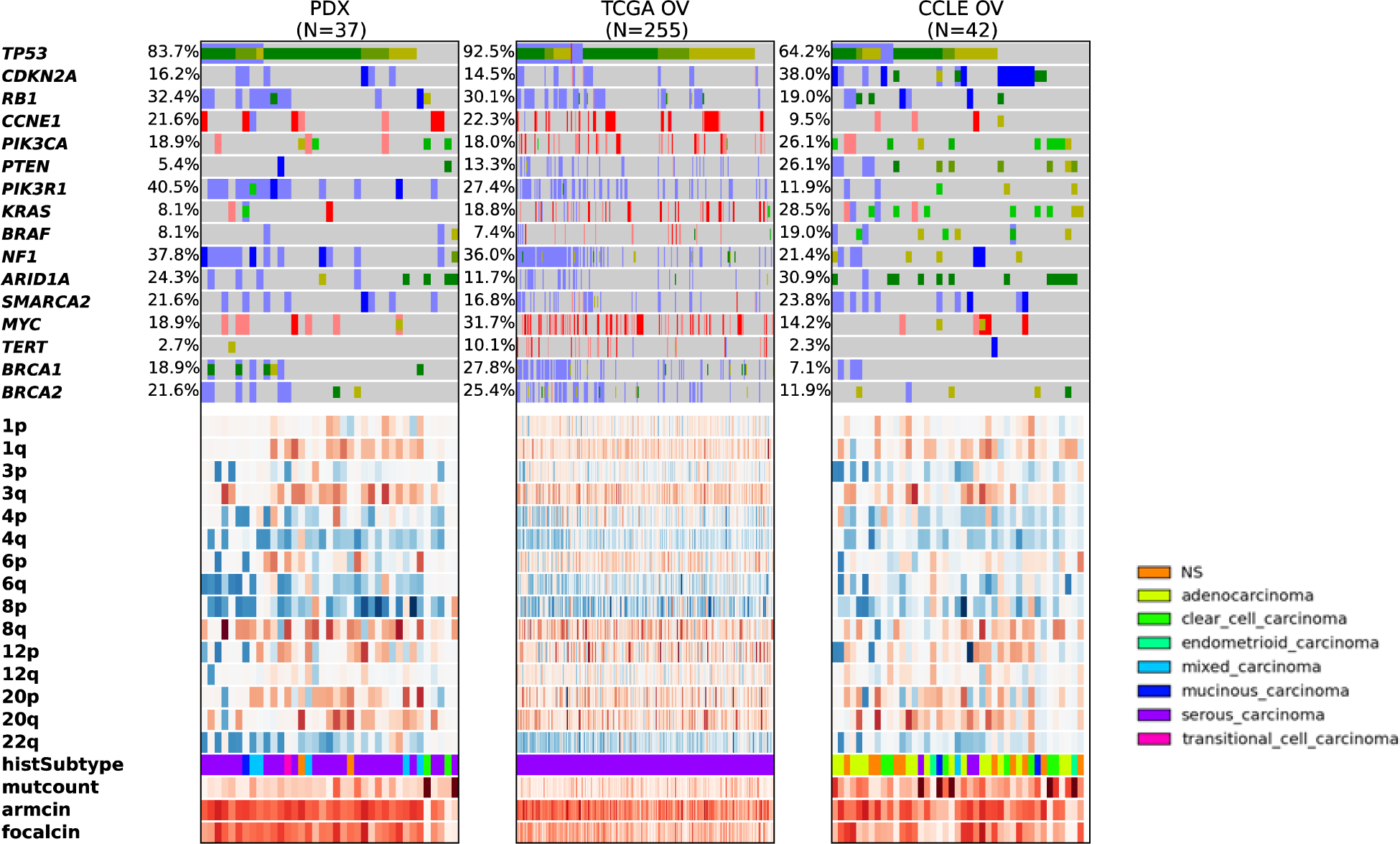
Genomic landscape analysis of epithelial ovarian carcinoma of PDXs, patient tumors (TCGA) and cell lines (CCLE). Parenthesis, number of models per indication. Legend is for histological subtypes. Blue, homozygous deletions; light blue, heterozygous deletion; salmon, amplification > 5 copies; red, amplification > 8 copies; bright green, known COSMIC gain-of-function mutations; dark green, truncating mutations / frameshift or known COSMIC loss-of-function; mustard, novel mutation.

**Supplementary Fig. S3.**
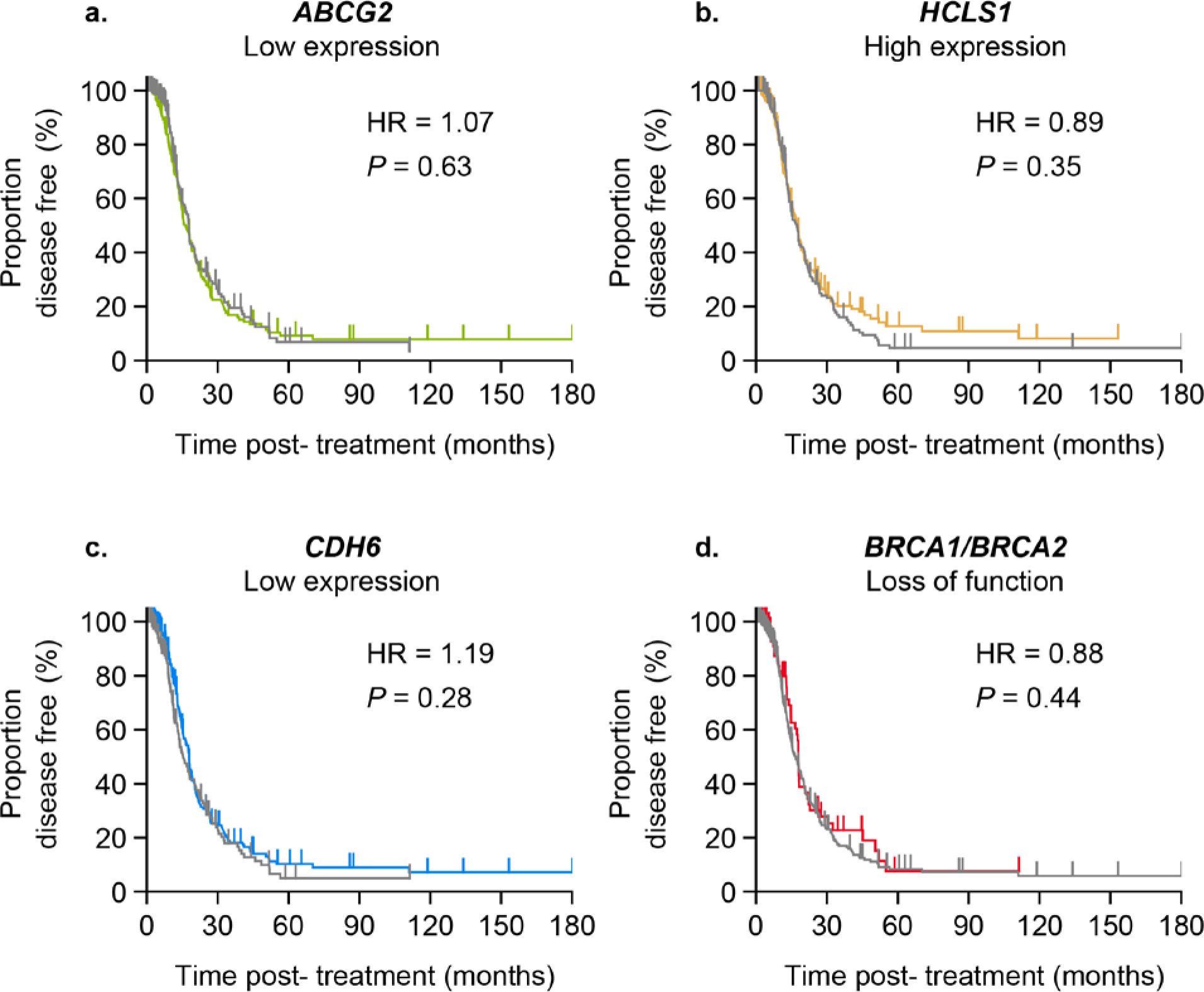
Survival analysis of primary ovarian tumor samples from TCGA. HR= hazard ratio, p-value calculated using Cox proportional hazards model. A) ABCG2 biomarker positive (green, Log2(ABCG2) < −1, n=155) versus biomarker negative (grey, Log2(ABCG2) >= −1, n=157). B) HCLS1 biomarker positive (gold, Log2(HCLS1) > 1.72, n=165) versus biomarker negative (grey, Log2(HCLS1) <= 1.72, n=147). C) CDH6 biomarker positive (blue, Log2(HCLS1) > 2.63, n=191) versus biomarker negative (grey, Log2(HCLS1) <= 2.63, n=121). D) BRCA1/BRCA2 biomarker positive (red, BRCA1/BRCA2 loss-of-function, n=61) versus biomarker negative (grey, BRCA1/BRCA2 wild-type, n=251).

**Supplementary Table 1**: Ovarian PDX subtype characteristics and take rate.

**Supplementary Table 2**: Ovarian PDX raw growth and sequencing data.

**Supplementary Table 3:** Ovarian PDX response category by therapy.

**Supplementary Table 4:** Mean tumor volume change difference between best monotherapy and each of 9 combinations. Hazard ratio for combination versus independent action simulation PFS.

**Supplementary Table 5:** Compound dose, dosing schedule, administration route, target and drug type.

## REFERENCES

1. Siegel, R. L., Miller, K. D. & Jemal A. Cancer statistics, 2015. CA Cancer J Clin 65, 5–29 (2015).

2. du Bois, A. et al. A randomized clinical trial of cisplatin/paclitaxel versus carboplatin/paclitaxel as first-line treatment of ovarian cancer. J. Natl. Cance. Inst. 95, 1320–1329 (2003).

3. Ozols, R. F. et al. Phase III trial of carboplatin and paclitaxel compared with cisplatin and paclitaxel in patients with optimally resected stage III ovarian cancer: a Gynecologic Oncology Group study. J. Clin. Oncol. 21, 3194–3200 (2003).

4. Coward, J. I., Middleton, K. & Murphy F. New perspectives on targeted therapy in ovarian cancer. Int J Womens Health 7, 189–203 (2015).

5. Yang, W. et al. Genomics of Drug Sensitivity in Cancer (GDSC): a resource for therapeutic biomarker discovery in cancer cells. Nucleic Acids Res 41, D955–D961 (2013).

6. Rees, M. G. et al. Correlating chemical sensitivity and basal gene expression reveals mechanism of action. Nat. Chem. Biol. 12, 109–116 (2016).

7. Yu, C. et al. High-throughput identification of genotype-specific cancer vulnerabilities in mixtures of barcoded tumor cell lines. Nat. Biotechnol. 34, 419–423 (2016).

8. Bertotti, A. et al. The genomic landscape of response to EGFR blockade in colorectal cancer. Nature 526, 263–267 (2015).

9. Gao, H. et al. High-throughput screening using patient-derived tumor xenografts to predict clinical trial drug response. Nat. Med. 21, 1318–1325 (2015).

10. Murphy, B. et al. Evaluation of Alternative In Vivo Drug Screening Methodology: A Single Mouse Analysis. Cancer Res. 76, 5798–5809 (2016).

11. Politi, K. & Herbst R. S. Lung Cancer in the Era of Precision Medicine. Clin Cancer Res 21, 2213–2220 (2015).

12. Korpal, M. et al. An F876L mutation in androgen receptor confers genetic and phenotypic resistance to MDV3100 (enzalutamide). Cancer Discov 3, 1030–1043 (2013).

13. Palmer, A. C. & Sorger P. K. Combination Cancer Therapy Can Confer Benefit via Patient-to-Patient Variability without Drug Additivity or Synergy. Cell 171, 1678–1691.e13 (2017).

14. Topp, M. D. et al. Molecular correlates of platinum response in human high-grade serous ovarian cancer patient-derived xenografts. Molecular Oncology 8, 656–668 (2014).

15. George, E. et al. A patient-derived-xenograft platform to study BRCA-deficient ovarian cancers. JCI Insight 2, (2017).

16. Dong, R. et al. Histologic and molecular analysis of patient derived xenografts of high-grade serous ovarian carcinoma. J Hematol Oncol 9, 92 (2016).

17. Cancer Genome Atlas Research Network. Integrated genomic analyses of ovarian carcinoma. Nature 474, 609–615 (2011).

18. Cheaib, B., Auguste, A. & Leary A. The PI3K/Akt/mTOR pathway in ovarian cancer: therapeutic opportunities and challenges. Chin J Cancer 34, 4–16 (2015).

19. Bialucha, C. U. et al. Discovery and Optimization of HKT288, a Cadherin-6-Targeting ADC for the Treatment of Ovarian and Renal Cancers. Cancer Discov 7, 1030–1045 (2017).

20. Gooijer, M. C. de et al. The impact of P-glycoprotein and breast cancer resistance protein on the brain pharmacokinetics and pharmacodynamics of a panel of MEK inhibitors. International Journal of Cancer 142, 381–391 (2018).

21. Gao, H. et al. High-throughput screening using patient-derived tumor xenografts to predict clinical trial drug response. Nat Med 21, 1318–1325 (2015).

22. Samareh, B. et al. Inhibition of the NAMPT-Triggered Deacetylation of HCLS1 Protein: A New Therapeutic Option in Chronic Myeloid Leukemia. Blood 126, 1001–1001 (2015).

23. Esteller, M. et al. Promoter Hypermethylation and BRCA1 Inactivation in Sporadic Breast and Ovarian Tumors. J Natl Cancer Inst 92, 564–569 (2000).

24. George, A., Kaye, S. & Banerjee S. Delivering widespread BRCA testing and PARP inhibition to patients with ovarian cancer. Nat Rev Clin Oncol 14, 284–296 (2017).

25. Wang, Z. et al. USP51 deubiquitylates H2AK13,15ub and regulates DNA damage response. Genes Dev. 30, 946–959 (2016).

26. Tsibulak, I., Zeimet, A. G. & Marth C. Hopes and failures in front-line ovarian cancer therapy. Critical Reviews in Oncology/Hematology 143, 14–19 (2019).

27. Townsend, E. C. et al. The Public Repository of Xenografts Enables Discovery and Randomized Phase II-like Trials in Mice. Cancer Cell 29, 574–586 (2016).

28. Tung, N. M. & Garber, J. E. BRCA 1/2 testing: therapeutic implications for breast cancer management. British Journal of Cancer 119, 141 (2018).

29. Moore, K. et al. Maintenance Olaparib in Patients with Newly Diagnosed Advanced Ovarian Cancer. N. Engl. J. Med. 379, 2495–2505 (2018).

30. Ferrarotto, R., Redman, M. W., Gandara, D. R., Herbst, R. S. & Papadimitrakopoulou V. A. Lung-MAP--framework, overview, and design principles. Chin Clin Oncol 4, 36 (2015).

31. Sethi, A. et al. The PDX Data Commons and Coordinating Center (PDCCC) for PDXNet. Cancer Res 78 (13 supplement), 1029 (2018).

