## Supplementary Table 1 for "Biomarker-guided treatment strategies for ovarian cancer identified from a heterogeneous panel of patient-derived tumor xenografts"

| Tumor subtype | # established models | # specimen received | % Take rate |
| --- | --- | --- | --- |
| Serous carcinoma | 34 | 154 | 22 |
| Clear cell carcinoma | 4 | 21 | 19 |
| Mucinous carcinoma | 1 | 12 | 8 |
| Transitional cell carcinoma/malignant Brenner tumor | 1 | 9 | 11 |
| Endometrioid carcinoma | 1 | 24 | 4 |
| Mixed carcinoma | 6 | 18 | 33 |
| Non-specified carcinoma | 2 | 9 | 22 |
