## Supplementary Table 4 for "Biomarker-guided treatment strategies for ovarian cancer identified from a heterogeneous panel of patient-derived tumor xenografts"

| **Combination Name** | **Mean tumor volume change difference between independent action and combination (%)** | **p-value** | **Hazard ratio for combination versus independent action simulation PFS** | **p-value** |
| --- | --- | --- | --- | --- |
| BKM120 + Binimetinib | -29 | 0.01 | 0.27 | 0.01 |
| BYL719 + LJM716 | -6 | 0.72 | 0.99 | 0.97 |
| CLR457 + LJM716 | -17 | 0.09 | 1.16 | 0.71 |
| BYL719 + Carboplatin + Paclitaxel | -1 | 0.96 | 0.70 | 0.68 |
| LEE011 + Carboplatin + Paclitaxel | 3 | 0.86 | 0.63 | 0.34 |
| BKM120 + Olaparib | 0 | 0.74 | 0.78 | 0.65 |
| CLR457 + Olaparib | 11 | 0.20 | 1.53 | 0.30 |
| CLR457 + Binimetinib | -1 | 0.85 | 0.80 | 0.62 |
| LEE011 + Binimetinib | -6 | 0.89 | 1.48 | 0.20 |
